# APALORD: An R-based tool for differential alternative polyadenylation analysis of long-read RNA-seq data

**DOI:** 10.1101/2025.06.11.658931

**Authors:** Zhiping Zhang, Heather Glatt-Deeley, Lehan Zou, Dongyuan Song, Pedro Miura

## Abstract

Alternative polyadenylation (APA) is a critical co/post-transcriptional process that enhances RNA isoform diversity, regulating mRNA stability, localization and translation in a spatiotemporal manner. Over the past decade, long-read (LR) RNA sequencing techniques have advanced rapidly, producing datasets that could offer insights into APA mechanisms. Here we introduce **APALORD** (Alternative Polyadenylation Analysis of LOng-ReaDs), an R-based analysis tool for APA analysis of LR RNA-seq data. Leveraging precise 3’ end information from 3’-primed LR RNA-seq data, APALORD identifies polyadenylation sites (PASs) and quantifies PAS usage (PAU) at individual sites for each sample. It conducts APA analysis at the gene level (using Kolmogorov-Smirnov test) and at the level of individual PAS (using DEXSeq) across sample conditions. APALORD was applied to direct RNA-seq data from human embryonic stem cells (hESCs) and hESC-derived neurons. PASs were identified with high accuracy and a transcriptome-wide 3’UTR lengthening trend was found, consistent with previous studies. APALORD analysis of PacBio cDNA data from human tissues confirmed a 3’UTR lengthening trend in cortex compared to liver. R2C2 libraries generated on the Nanopore platform were analyzed with APALORD and it revealed APA change associated with polysome fractions in human neural progenitor cells. In summary, APALORD offers a comprehensive framework for differential APA analysis using LR RNA-Seq data, empowering researchers to investigate 3’ end dynamics across diverse biological contexts.

## Introduction

^1^During mRNA processing, the cleavage and polyadenylation (CPA) machinery recognizes specific cis-acting sequence elements surrounding the cleavage site, to collectively define a polyadenylation site (PAS)^1,2,3^. The core element is the hexamer sequence AAUAAA (or its variants), bound by the CPA specificity factor (CPSF)^4,5^. Additional elements include the UGUA motif (upstream sequence element, USE) bound by cleavage factor I (CFI)^6,7^, and a GU/U-rich downstream sequence element (DSE) bound by cleavage stimulation factor (CSTF)^8,9^. Alternative polyadenylation (APA) is a tightly regulated co/post-transcriptional process that generates alternative 3’ends for genes^10^. Most human genes generate more than one 3’UTR mRNA isoform due to APA^11^. For broadly expressed genes, tissue specific patterns of APA are found across phyla, with distal PAS selection preferred in the brain and proximal PAS selection preferred in testis^12,13^. Cell stress, development, and differentiation regulate APA patterns^14,15^. APA can produce mRNAs with alternative C-terminal coding potential, but most commonly affects 3’UTR sequence content (last exon tandem APA) that can alter mRNA stability, localization, and translational control. Roles for alternative 3’UTR mRNA isoforms include nervous system development, axon growth, mating behavior^16,17,18,19^. APA patterns are disrupted in several diseases, including glioma^20^ and APA quantitative trait loci contributes to disease heritability^21^.

While transcriptome-wide APA quantification is feasible with conventional Illumina-based short-read RNA sequencing, it faces trade-offs between gene body coverage and 3’ end accuracy or depth^22^. APA analysis using standard, non–3’ end tag-specific short-read library preparations often lacks precise information about 3’ ends. In contrast, 3’ end tag-specific short-read library preparations improve both the accuracy and depth of 3’ end measurements. However, their poor gene body coverage makes it less suitable for widespread application and limited read lengths (typically 50–300 bp) can lead to ambiguous alignment to specific genes or isoforms^23^. Consequently, short-read APA analysis typically relies on well-annotated PAS references to estimate 3’UTR isoform usage or requires predefined regions to detect APA changes across conditions^24,25,26^. Poor PAS annotation and uneven coverage in short-read data frequently distort gene-level APA analyses.

The advent of long-read (LR) RNA sequencing a decade ago initially aided the discovery of rare isoforms poorly resolved by short-read methods, though its low depth and high error rate limited broader applications^27,28^. Recent advances in high-throughput LR RNA-seq have significantly improved its accuracy and depth, establishing it as a powerful tool for RNA profiling^29,30^, offering reduced ambiguity and comparable depth to traditional short-read RNA-seq techniques^31^. LR sequencing captures full-length RNA or cDNA sequences, enabling precise gene and isoform assignment as well as accurate quantification^32,33,34^. Moreover, LR data exhibit higher 3’ end coverage and more consistent depth over longer regions (> 1 kb) compared to short-read data, facilitating data-driven PAS detection and accurate quantification of 3’UTR isoforms. Direct RNA-seq, which sequences the full length of the native RNA strand from the end of the polyadenosine tail without any added chemical modifications, offers the most accurate 3’UTR information^35,36^. Direct RNA-seq data avoids artifacts stemming from PCR amplification and internal poly(A) priming, which plague the aforementioned APA analysis. Currently, tools for transcriptome-wide APA analysis of LR RNA-seq are scarce.

Here, we introduce APALORD (Alternative Polyadenylation Analysis of LOng ReaDs), an R-based tool for quantifying differentially regulated APA events from long-read datasets. APALORD performs statistical analyses at individual PAS level to identify differentially used sites and detects trends in 3’UTR shortening/lengthening at the gene level across samples. We profiled hESC samples during neural differentiation to confirm a global trend for 3’UTR lengthening. We also demonstrate the utility of APALORD on cDNA datasets from PacBio and Oxford Nanopore platforms. With the growing adoption of long-read RNA-seq, we anticipate APALORD will become a widely used tool for investigating 3’ end transcript dynamics across diverse biological systems.

## Results

### APALORD computational workflow

The APALORD R package analyzes LR RNA-seq data to provide statistical analysis of differential expression at the individual PAS level and a readout of 3’UTR shortening/lengthening trends at the gene level. LR RNA-seq reads are first aligned to the host genome using *minimap2*^376^, followed by assignment to transcribed genes using IsoQuant^33^. The output data from IsoQuant is then loaded into R for APA analysis with APALORD (Fig.1). APALORD requires the same GTF reference file used by minimap2 and IsoQuant to extract general gene information—such as gene name, stop codon position, and strand—but does not rely on existing 3’UTR annotations. Using precise 3’ end positions identified by IsoQuant from aligned reads, APALORD annotates PASs, defined as peaks where most reads from a given PAS cluster within a short window (≤ 20 nt) around the peak. Users can apply a customizable cutoff to retain only highly expressed PASs of genes with multiple 3’ PASs (e.g., percent of PAS usage (PAU) ≥ 5%, with a default of 1%) for downstream analysis.

**Fig. 1:**
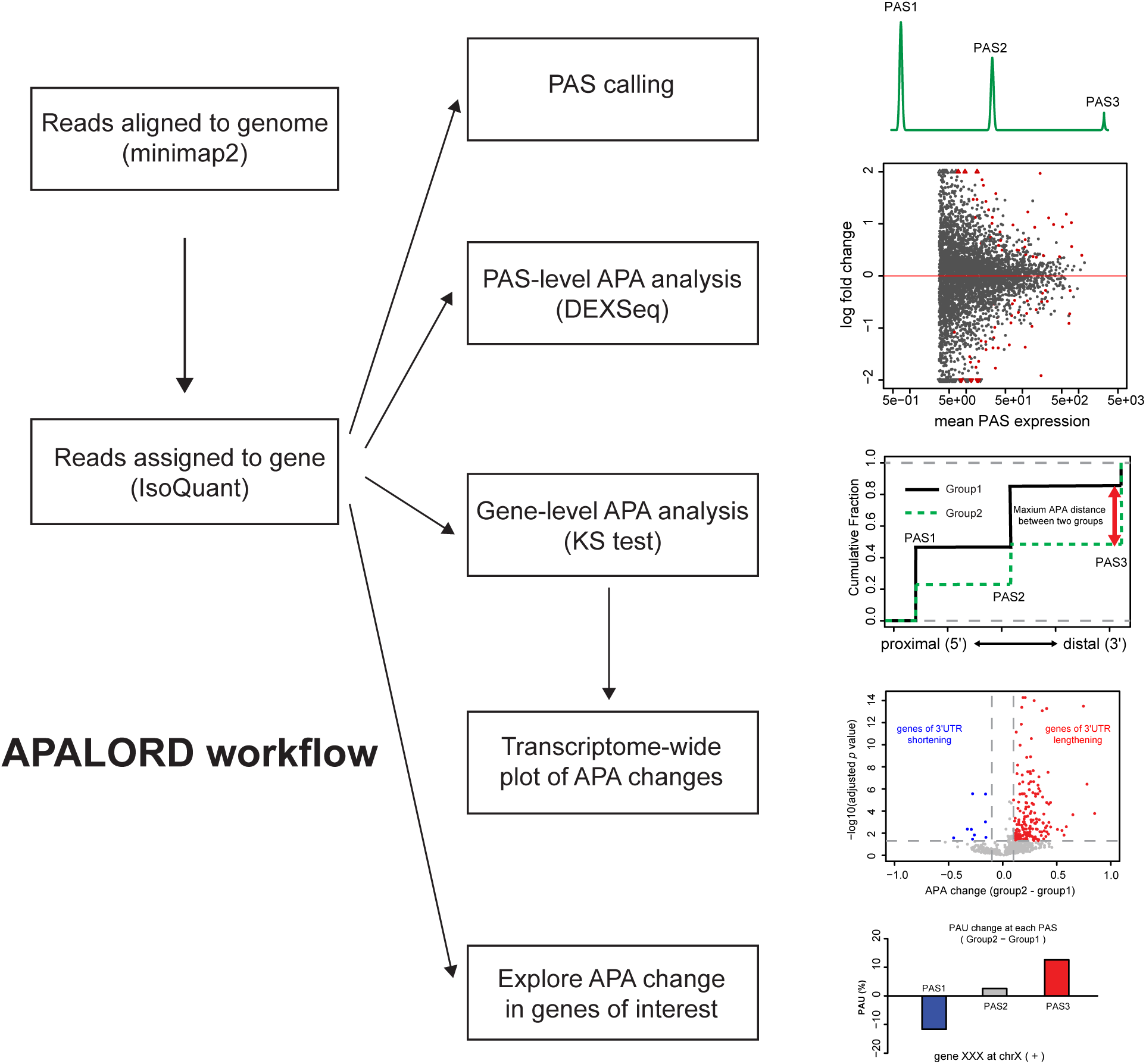
APALORD workflow. APALORD workflow. Read sequences are first aligned to the reference genome using minimap2, producing a BAM file. IsoQuant then assigns these aligned reads to unique genes. APALORD begins with read assignment information generated by IsoQuant. APALORD offers a suite of analyses, including PAS calling, PAS-level APA analysis via PAS usage (PAU) quantification, gene-level APA analysis with visualization across the transcriptome and single-gene APA change visualization.

The APALORD workflow (Fig. 1) includes analysis at the individual PAS level where it employs DEXSeq^38^, originally designed for differential exon usage, to identify differentially used PAS between two conditions (e.g., control vs. experimental), each with at least two replicates. This analysis detects PASs with significant usage differences, regardless of their relative position within the host gene. To assess general 3’UTR lengthening (i.e., distal PAS preferential selection) or shortening (i.e., proximal PAS preferential selection), APALORD applies the non-parametric Kolmogorov-Smirnov (KS) test to compare the distributions of 3’ end positions along the gene body between conditions. The KS test yields a *p*-value and a maximum difference (ranging from -1 to 1) between cumulative distribution functions (CDFs), serving as an indicator of gene-level APA changes. These data are visualized transcriptome-wide by volcano plot (APA change vs. adjusted *p*-value). To explore the origin of APA change in a gene of their interest, 3’ end distribution of all the reads assigned to the gene is plotted per condition and PAU change between conditions at all the PASs detected within the gene are visualized by bar plot.

### Applying APALORD to call PASs using hESC neural differentiation direct RNA-seq **data**

To demonstrate the performance of APALORD, we differentiated human NGN2-inducible H9 embryonic stem cells (hESC) into neurons^39^ and conducted direct RNA-seq on the Nanopore platform. Samples were collected from undifferentiated cells (D0) and neurons after 7 days NGN2-induced neural differentiation (D7) (Fig.2a). 15,297,408 reads from the hESCs were assigned uniquely to 25,066 genes, and 8,977,059 reads from the neurons were assigned to 24,236 genes (28,290 genes together). From this dataset (triplicates for both D0 and D7 condition), APALORD identified 54,674 PASs (overall PAU ≥1%), of which 39,817 (72.8%) had sufficient depth for PAS usage (PAU) calculation. Each PAS annotation was collapsed to the nearby nucleotide coordinate with the most read depth (within 20 nt). Comparing these PASs to the annotated PASs in PolyA_DB^40,41^, we found that 26,046 (65.4%) matched annotated PASs within the same host gene. Additionally, 7,358 (18.5%) PASs were located within 20 nt of a known PAS, with 31,821 (95.3%) of these within 10 nt (Fig.2b). The distribution of distances between APALORD-detected PAS peaks and nearby PolyA_DB PASs confirmed high accuracy of APALORD in pinpointing PAS peak positions (Fig. 2c, *p* < 2.2e-16, Chi-square test). The remaining 6,413 PASs (16.1%) were classified as novel (Fig. 2b).

**Fig. 2:**
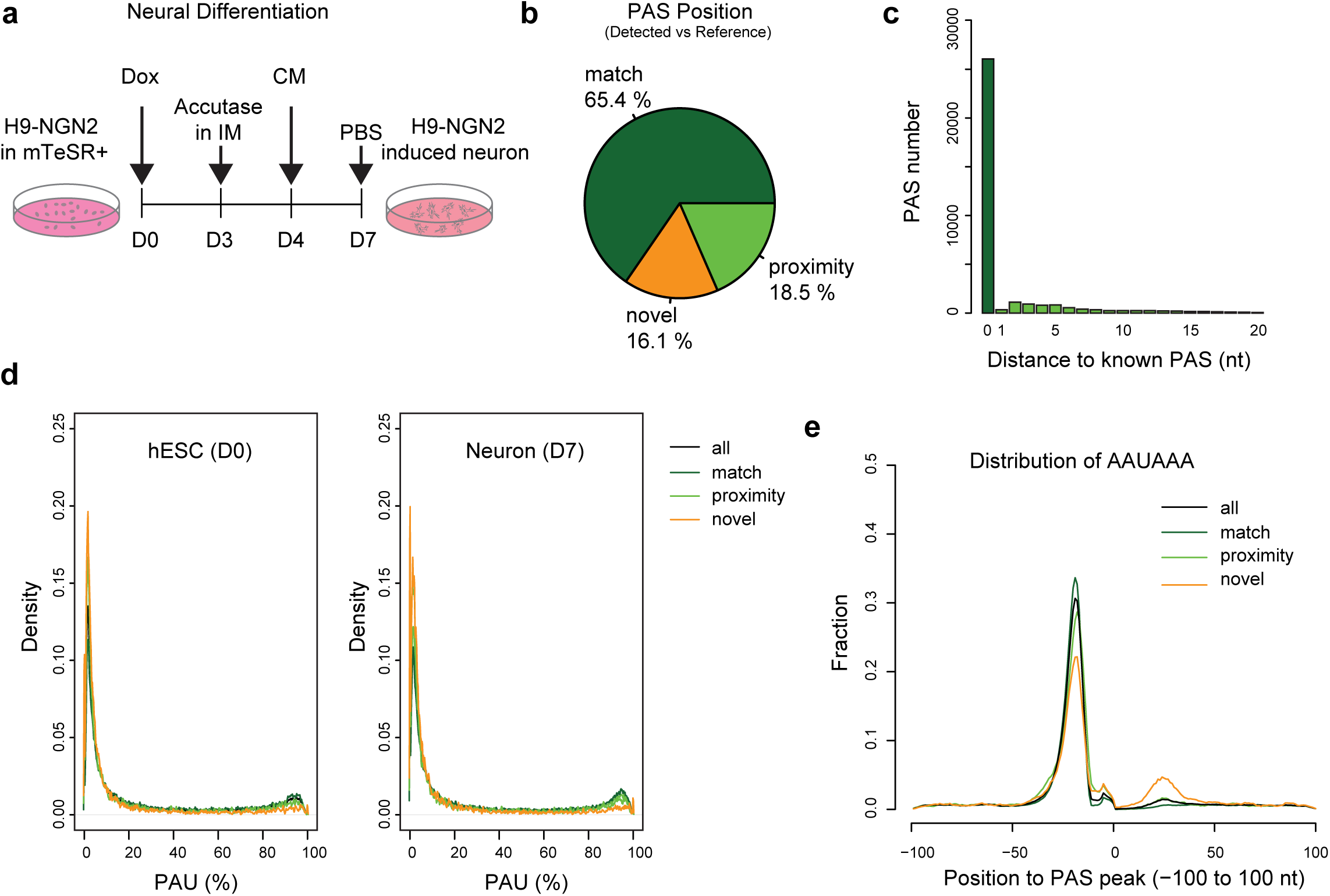
Performance of APALORD in PAS calling from direct RNA-seq of hESCs and hESC-derived neurons. **a** Schematic of neural differentiation from NGN2-inducible H9 ESCs (D0) to induced neurons (D7). Dox: doxycycline. IM: induction medium. CM, cortical neuron medium. **b** PASs identified by APALORD from direct RNA-seq data of hESCs and hESC-derived neurons. match: exact alignment with annotated PASs in the PolyA_DB reference; proximity: within the distance of 20-nt to an annotated PAS; novel: outside of match or proximity criteria. **c** Distance distribution of match and proximity PASs to annotated PASs. **d** Density plot of PAU for D0 and D7 samples across all, match, proximity and novel PASs. **e** Proportion of PASs with the canonical AAUAAA hexamer within the sequence surrounding the peak cleavage site (-100 nt to +100 nt). Categories of PASs as defined in **b**.

We observed that novel PASs in both D0 and D7 samples exhibited lower expression compared to known PASs or PASs in proximity (Fig. 2d). This aligns with our expectation that APALORD’s sensitivity, driven by precise 3’ end detection, enables identification of lowly expressed PASs. We extracted sequences flanking PAS peaks (-100 to +100 nt) and found that novel PASs were less likely to contain the canonical AAUAAA motif upstream compared to annotated PASs but had a higher probability of featuring an AAUAAA motif in the downstream 10 to 30 nt sequence (Fig. 2e). Among the 11,341 genes with identified PASs, 8,963 genes (79%) utilized 2 or more PASs, with a median of 4 PASs per gene. Notably, more than half of expressed genes (6,560) used 3 or more PASs (Supplementary Fig.1b). These observations are consistent with previous reports^42,41,11^. Among novel PASs, 46.7% resided in known 3’UTR exons, followed by 29.6.1% in introns and 16.4% in coding sequence (CDS) exons (Supplementary Fig. 1c).

### PAS-level and gene-level APA analysis during neural differentiation

We performed single-PAS level statistical analysis between hESC and hESC-derived neuron samples (Fig.3a) using a matrix of read counts per PAS per sample. This revealed 3544 significantly regulated PASs (fold change (FC) >1.5 and adjusted *p* < 0.05) across 2383 genes, with 1771 up-regulated and 1773 down-regulated in neurons (Fig.3b). Employing a Kolmogorov-Smirnov (KS) test to compare the 3’ end distribution between neuron and hESC samples, we called genes with a trend to use more proximal (5’) or distal (3’) PASs (Fig.3c). This gene-level APA analysis detected 2,138 genes with a 3’UTR lengthening trend and 158 with a shortening trend (adjusted *p* < 0.05, APA change > 0.1 or < -0.1, Fig. 3d). APALORD categorizes genes subjected to APA regulation into two groups. The first, “Last-exon tandem APA”, identifies APA genes where alternative PASs reside solely in the last exon, thus reflecting 3’UTR APA. The second, defined as “mixed APA”, encompasses genes expressing both intronic and last-exon PASs). We found such classifications necessary because of the many instances where a given gene was regulated by both tandem APA and alternative last exon APA. For the mixed APA category, APA shifts may involve intronic PASs, last-exon 3’UTR PASs, or both. Among genes exhibiting 3’UTR lengthening from hESCs to neurons, last-exon tandem APA regulation events outnumbered mixed APA regulation events (Supplementary Fig. 2a). Gene Ontology (GO) analysis of biological processes highlighted predominantly neural-related functions for the differential APA events (Supplementary Fig. 2b).

**Fig. 3:**
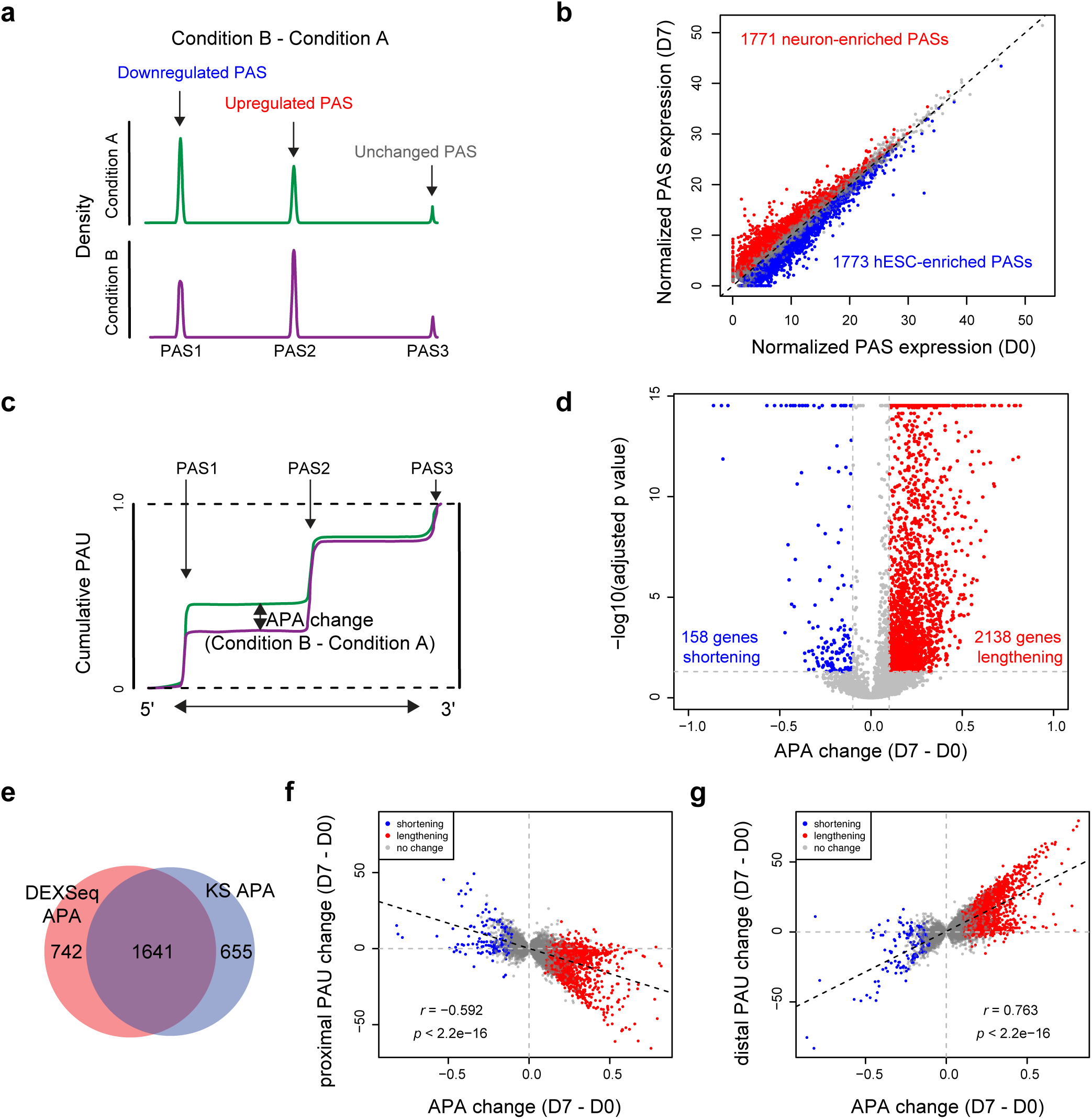
APALORD analysis of APA changes during hESC neural differentiation. **a** APALORD quantifies reads terminating at each PAS in a target gene and identifies regulated PAS events between two conditions using DEXSeq. **b** Normalized PAS expression of individual PASs in hESC (D0, n=3) and hESC-derived neuron (D7, n=3) samples. Red: neuron-enriched PASs (D7 > D0, adjusted *p* < 0.05). Blue: hESC-enriched PASs (D7 < D0, adjusted *p* < 0.05). **c** APALORD applies the Kolmogorov-Smirnov (KS) test to detect the maximum difference of cumulative PAUs between two conditions and identify significantly regulated APA genes. **d** Volcano plot illustrating transcriptome-wide APA changes during hESC neural differentiation. Red: genes with lengthened 3’UTR in neurons (APA change (D7 – D0) > 0.1, adjusted *p* < 0.05). Blue: genes expressing shortened 3’UTR in neurons (APA change (D7 – D0) < -0.1, adjusted *p* < 0.05). **e** Venn diagram showing overlap between neural differentiation-regulated APA genes identified by PAS-level APA analysis using DEXSeq and gene-level APA analysis using KS test. DEXSeq APA: | log2(FC(D7/D0))| > log2(1.5), adjusted *p* < 0.05, at least one significant PAS event in the genes. KS APA: |PAU change (D7 – D0) | > 0.1, adjusted *p* < 0.05. **f** Correlation between KS test-reported APA change and PAU change at the proximal PAS in single genes during neural differentiation. Pearson’s correlation, *r* = -0.592, *p* < 2.2e-16. **g** Correlation between KS test-reported APA change and PAU change at the distal PAS in single genes during neural differentiation. Pearson’s correlation, *r* = 0.763, *p* < 2.2e-16. Red and blue points in **f** and **g** represent significant APA genes from **d**.

Comparing APA regulated genes identified by PAS-level (by DEXSeq) and gene-level (by KS test) analyses, 1,641 genes overlapped (Fig. 3e), representing 68.9% of DEXSeq APA and 71.5% of KS APA events. Preferred usage of more proximal or distal PAS has been pointed out as 3’UTR lengthening or shortening trends when comparing conditions^25,43^. We found that gene-level APA changes correlated significantly and negatively with proximal PAU changes (*r* = - 0.592, *p* < 2.2e-16) but positively with distal PAU changes (*r* = 0.763, *p* < 2.2e-16) during neural differentiation (Fig. 3f and 3g). This aligns with prior findings that distal PASs are preferentially used in the nervous system and neurons^44,12,45^.

### APA regulation trends for individual genes revealed by APALORD

APALORD provides visualization tools of APA regulation events for individual genes. This is especially helpful given the complexity of APA regulation, which can often include 3 or more different PASs per gene. The *NAV1* gene (Neuron Navigator 1) plays a role in axon outgrowth and migration^46^. In our direct RNA-seq data, APALORD identified that *NAV1* undergoes cleavage and polyadenylation at 4 tandem 3’UTR PASs. *NAV1* was classified as exhibiting significant 3’UTR lengthening during neural differentiation, with PAS1 and PAS4 identified as regulated PAS events (Fig.4a-d). Specifically, PAS1 usage decreased from 30.1% to 11.8%, while PAS4 usage increased from 8.6% to 35.3%. *SYNGR1*, which encodes a protein associated with presynaptic vesicles in neuronal cells^47^, utilizes 4 PASs that span multiple exons. The primary PAU switch occurred between intronic PAS1 and distal PAS4, with PAS1 usage dropping from 70.4% to 17.8% and PAS4 usage rising from 10.4% to 69.6% (Fig.4e-h).

**Fig. 4:**
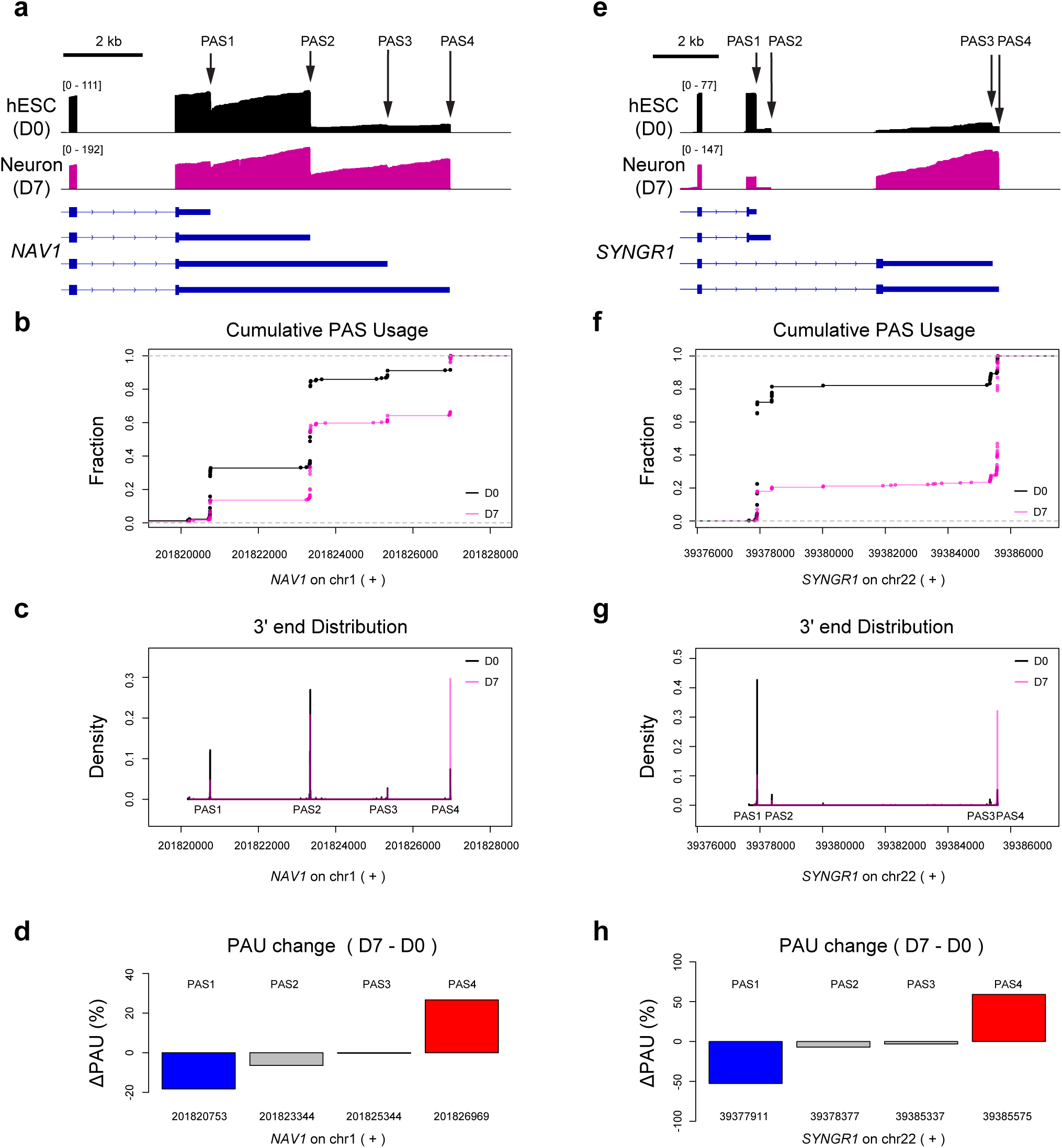
APALORD visualization of differential APA during hESC neural differentiation for *NAV1* and *SYNGR1* genes. **a**-**d** Analysis of the last exon tandem APA gene *NAV1* in hESC (D0) and hESC-derived neuron (D7) samples: direct RNA-seq coverage tracks (**a**), cumulative PAU curves (**b**), density plots of 3’ ends (**c**) and PAU change (D7 – D0) (**d**) at individual PASs identified by APALORD. **e**-**h** Analysis of the mixed APA gene *SYNGR1* in hESC (D0) and hESC-derived neuron (D7) samples: direct RNA-seq coverage tracks (**e**), cumulative PAU curves (**f**), density plots of 3’ ends (**g**) and PAU change (D7 – D0) (**h**) at individual PASs identified by APALORD.

### Sequence features of neural differentiation regulated PASs

To investigate motifs and nucleotide distribution patterns potentially regulating alternative PAS usage, we extracted sequences (-100 to +100 nt) flanking the cleavage sites of all reads in the D0 and D7 datasets. Nucleotide distributions revealed upstream A- and U-rich regions and downstream U-rich elements, consistent across both conditions (D0 and D7; Fig. 5a, 5b, and 5c). We further analyzed sequences around 11,102 PAS peaks with sufficient depth (>10 reads across all samples), including 1,308 neuron-enriched PASs (PAU_D7 – PAU_D0 >10) and 800 hESC-enriched PASs (PAU_D0 – PAU_D7 >10). Neuron-enriched PASs exhibited a pronounced upstream A-rich and downstream U-rich pattern compared to other groups (Fig. 5d). Given that the A-rich region overlaps the typical location of the canonical AAUAAA poly(A) signal (-35 to - 10 nt), we assessed AAUAAA motif incidence around PAS peaks and found a higher prevalence in the upstream sequences of neuron-enriched PASs (Fig. 5e and 5f). Similarly, the UGUA motif, an upstream sequence element, showed increased AAUAAA incidence in the -100 to -30 nt region of neuron-enriched PASs (Fig. 5g). These findings suggest that genes in neurons tend toward expression of stronger PASs compared to hESCs.

**Fig. 5:**
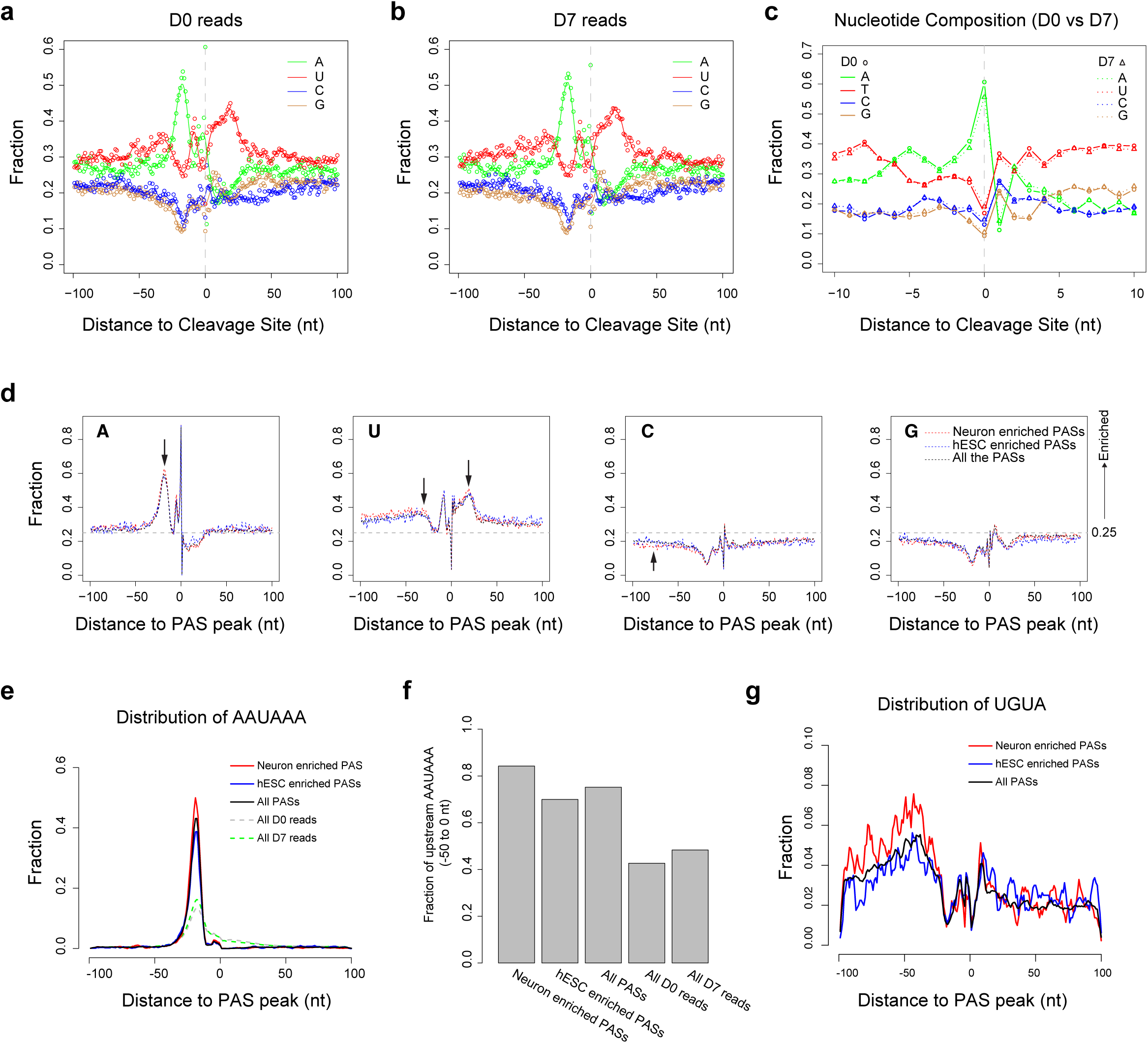
Nucleotide composition around 3’ cleavage sites in hESC and hESC-derived neuron samples. **a** Nucleotide composition from -100nt to +100nt around 3’ cleavage sites in hESC (D0) transcripts. **b** Nucleotide composition from -100nt to +100nt around 3’ cleavage sites in hESC-derived neuron (D7) transcripts. Points in **a** and **b** represent the fraction of each nucleotide (A, U, C, G) at each position; lines in **a** and **b** show the smoothed nucleotide fraction trends. **c** Nucleotide composition from -10nt to +10nt around 3’ cleavage sites in hESC (D0) and hESC-derived neuron (D7) transcripts. **d** Nucleotide composition from -100nt to +100nt around PAS peaks in hESC (D0) and hESC-derived neuron (D7) samples. Red: neuron-enriched PASs (PAU (D7 – D0) > 0.1, read counts > 20 per sample). Blue: hESC-enriched PASs (PAU (D7 – D0) < -0.1, read counts > 20 per sample). Black: all expressed PASs (read counts > 20 per sample). **e** Distribution of the AAUAAA motif from -100nt to +100nt around PAS peaks (red: neuron-enriched; blue: hESC-enriched; black: all expressed PASs) or reads 3’ cleavage sites (gray: all D0 transcripts; green: all D7 transcripts). **f** Proportion of PASs with the AAUAAA motif in the 50nt upstream of PAS peaks or reads 3’ cleavage sites. **g** Distribution of the UGUA motif from -100nt to +100nt around PAS peaks (red: neuron-enriched; blue: hESC-enriched; black: all expressed PASs).

### Applying APALORD to PacBio RNA-seq data

We applied APALORD to cDNA libraries sequenced on the PacBio LR sequencing platform^48^. Previous APA analyses identified a trend for 3’UTR lengthening in several brain regions compared to liver in both mouse and human^13^. We analyzed three dorsolateral prefrontal cortex libraries and two right liver lobe libraries, comprising 3,988,909 reads (1,145,512 from liver and 2,843,396 from cortex). APALORD identified 12,476 PASs across 2,975 genes with sufficient read depth. 392 (3.1%) of these PASs matched annotated PASs in the PolyA_DB reference, while 6773 (54.3%) were located in the proximity of known PASs in database (Supplementary Fig.4a).

Given that 42.6% of PASs were novel, we investigated whether internal priming, which often generates 3’ end artifacts, contributed to this observation. We labeled PASs surrounded by A-rich stretches (>6As in 10nt pre- or post-PAS peak sequences) as potential internal priming artifacts in APALORD. This filtering excluded 2,372 (1625 were novel) PASs due to suspected internal priming at the PAS peaks. Among the remaining “high-confidence” PASs, 389 (3.8%) matched annotated PASs and 6,030 (59.7%) were in the proximity of annotated PASs (Fig.6a). 6,047 (94.2%) PASs were found within 10nt of known PASs (Fig.6b). The 3,685 (36.5%) novel PASs were less likely to be supported by the upstream canonical AAUAAA hexamer when compared with PASs either matching or near known PASs in the reference (Fig.6c). Filtering for internal priming effectively removed PASs which might be artifacts caused by internal priming and lack support from a canonical upstream poly(A) hexamer (Supplementary Fig.4b). Gene-level APA analysis revealed 369 genes with 3’UTR shortening and 798 genes with lengthening in cortex compared to liver (Fig.6d). This significant trend toward 3’UTR lengthening (*p* < 0.00001, Fisher’s exact test) was consistent with previous findings^13^. Although the transcriptome-wide APA trend was identified by APALORD, the 3’ end position accuracy was lower in PacBio cDNA data compared to direct RNA-seq data. Internal priming-associated PASs accounted for 19% of expressed PASs but were present in 1,415 (59.4%) of 2,382 APA-analyzed genes, particularly APA regulated genes (Fig.6e). Of 1,167 APA regulated genes, only 431 (36.9%) had significantly regulated PASs. In contrast, 84.2% of DEXSeq-detected APA genes were also identified by KS test-based gene-level APA analysis (fold change > 1.5 or < 2/3, adjust *p* < 0.05; Fig.6f). As PASs in DEXSeq analysis were filtered to exclude internal priming artifacts, the discrepancy between these approaches suggests there is a higher degree of likely false positives for significant events identified by gene-level APA analysis due to internal priming artifacts.

**Fig. 6:**
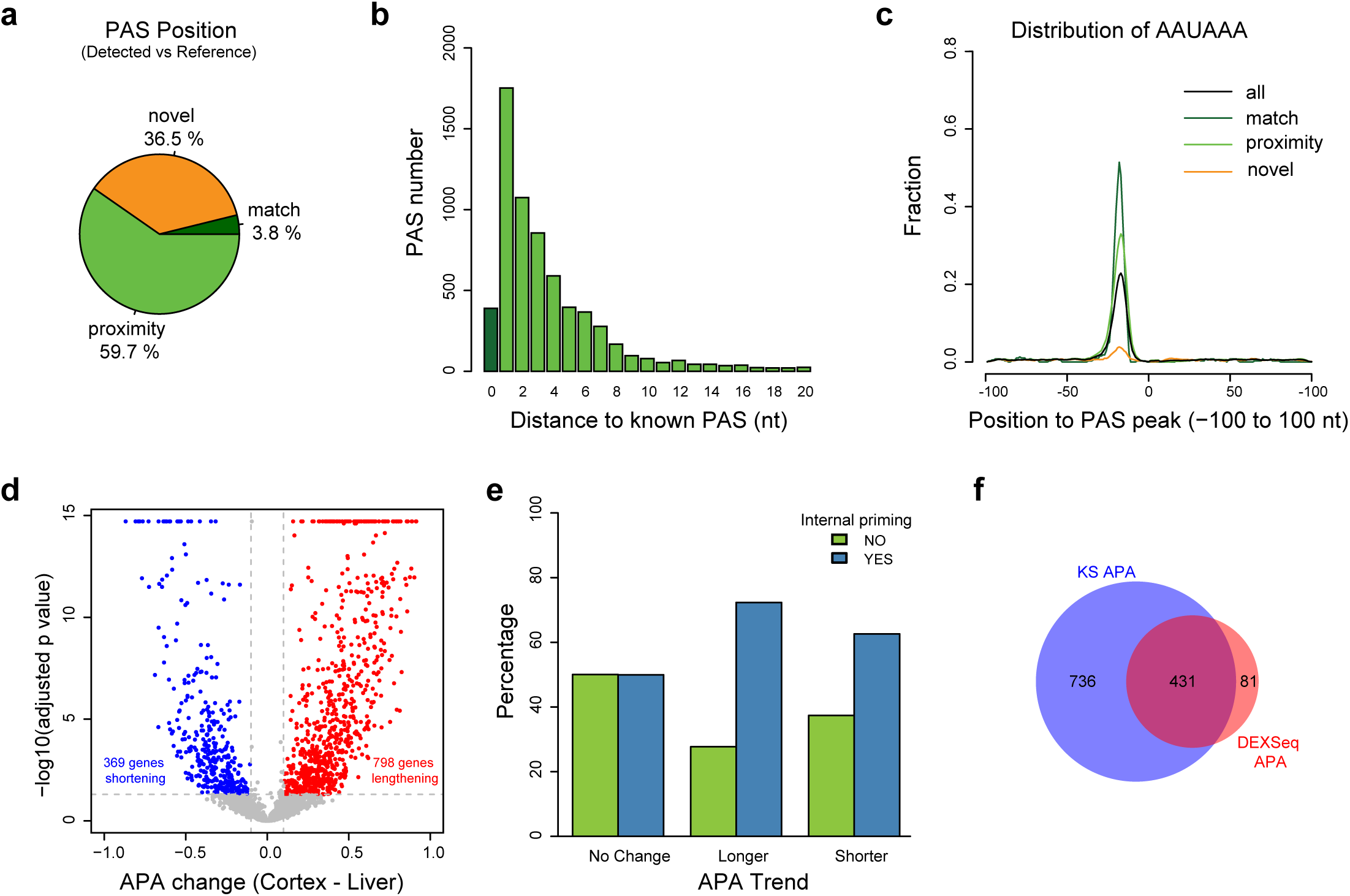
APALORD performance in analyzing PacBio cDNA data from human cortex and liver. **a** PASs identified by APALORD after excluding potential internal priming artifacts from PacBio cDNA data of human cortex and liver, categorized as: match (aligned with annotated PASs in the PolyA_DB reference), proximity (within 20 nt of an annotated PAS), novel (outside match and proximity criteria). **b** Distance distribution of match and proximity PASs to annotated PASs. **c** Proportion of PASs with the canonical AAUAAA hexamer within the sequence surrounding the peak cleavage site (-100 nt to +100 nt). Categories of PASs as defined in **a**. **d** Volcano plot illustrating transcriptome-wide APA changes between liver and cortex. Red: with lengthened 3’UTR in cortex (APA change (Cortex - Liver) > 0.1, adjusted *p* < 0.05). Blue: genes with shortened 3’UTR in cortex (APA change (Cortex - Liver) < -0.1, adjusted *p* < 0.05). **e** Counts of APA genes identified by APALORD between cortex and liver, categorized by presence of potential internal priming artifacts and 3’UTR change (lengthening, shortening or no change) as defined in **d**. **f** Venn diagram showing overlap between DEXSeq identified significant APA genes and KS test identified significant APA genes. DEXSeq APA: |log2(FC(Cortex/Liver)| > log2(1.5), adjusted *p* < 0.05, at least one significant PAS event in the genes. KS APA: |PAU change (Cortex – Liver)| > 0.1, adjusted *p* < 0.05.

### Polysome profiling using LR reveals translation activity associated APA changes

APA influences translational efficiency via changes in alternative 3’UTR sequence usage^10,49,50^. To explore this relationship, we analyzed long-read RNA-seq data from polysome fractions of hESC and hESC-derived neural progenitor cell (NPC) samples^51^. Cytosol extracts from ESC and NPC samples were fractionated on sucrose gradients to isolate monosome (associated with single ribosome), light polysome (2-4 ribosomes), and heavy polysome (≥5 ribosomes) in addition to whole cytosolic mRNA fraction, followed by library preparation with the R2C2 method and sequencing on the Nanopore platform^52^. Using APALORD, we confirmed a global trend of 3’UTR lengthening in all NPC fractions compared to hESC fractions (Supplementary Fig.4). The cytosol RNA fraction exhibited the fewest APA changes during neural differentiation from ESCs to NPCs, suggesting that mRNA from APA-regulated genes was enriched in polysome fractions. By analyzing neural differentiation-regulated APA genes from these four fractions, we noticed that the NPC lengthening genes were most abundant in the light polysome fraction (Fig.7a and Supplementary Fig.4). The NPC shortening genes showed fraction-specific patterns, with few genes consistently shortened across multiple fractions (Fig.7b).

**Fig. 7:**
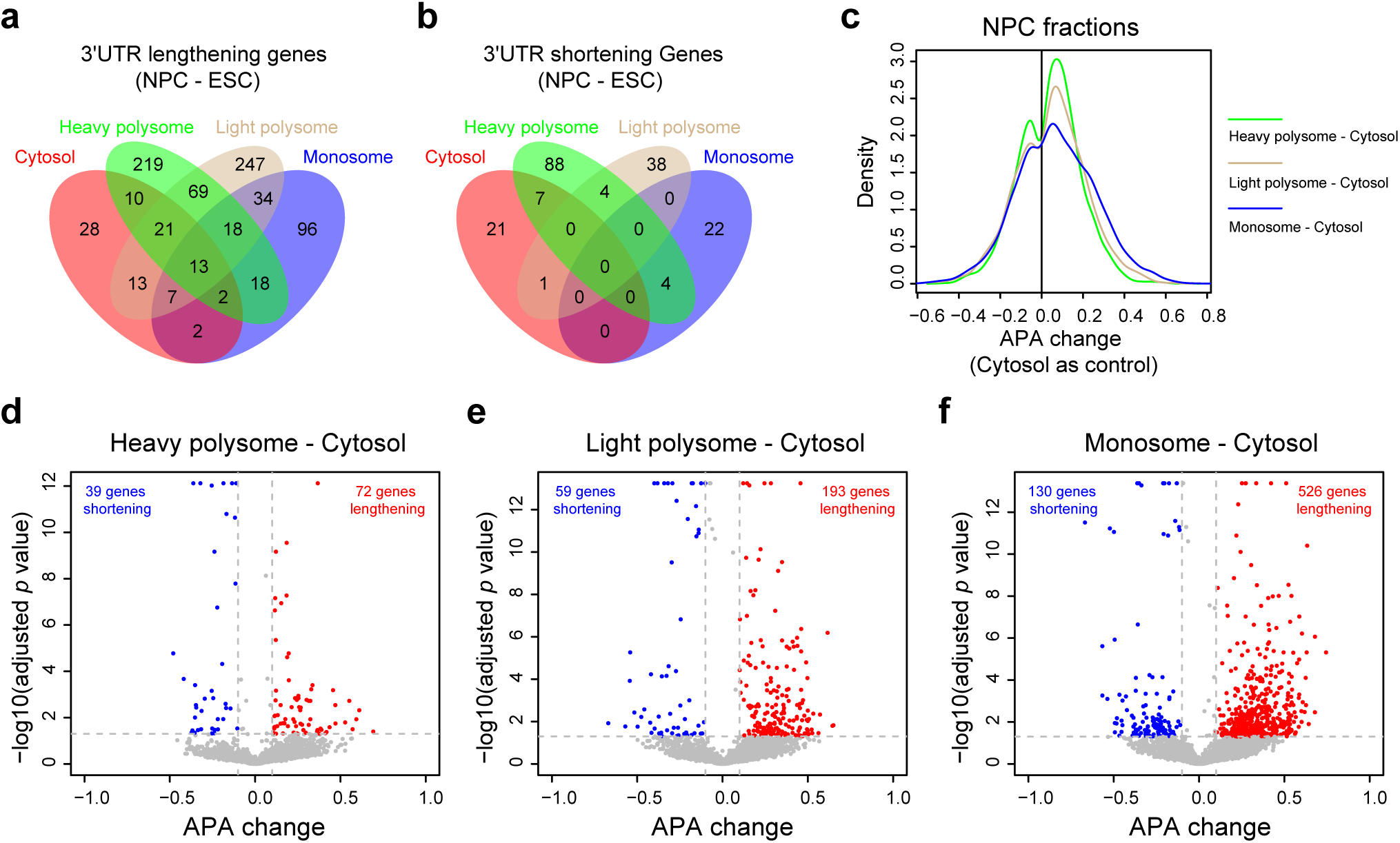
APALORD applied to polysome profiling using Nanopore R2C2 LR RNA-seq reveals translation activity associated APA changes. **a** Venn diagram showing the overlap of 3’UTR lengthening genes in NPCs compared to ESCs across fractions identified by APALORD from Nanopore R2C2 LR RNA-seq data, as defined in Supplementary Fig.4. **b** Venn diagram showing overlap of 3’UTR shortening genes in NPCs compared to ESCs across fractions identified by APALORD from Nanopore R2C2 LR RNA-seq data, as defined in Supplementary Fig.4. **c** APA change in NPC polysome fractions compared to NPC cytosol samples. **d-f** Transcriptome-wide APA changes quantified by APALORD in NPC heavy polysome (**d**), light polysome (**e**) and monosome (**f**) fractions compared to cytosol.

To investigate whether 3’UTR lengthening in NPCs is polysome fraction-dependent, we compared polysome fractions to the cytosolic fraction in NPCs. Gene-level APA changes increased progressively from heavy to light polysome fractions, peaking in the monosome fraction (Fig.7c). Consistently, genes with 3’UTR lengthening were most enriched in the monosome fraction, followed by the light polysome fraction, and least in the heavy polysome fraction (Fig.7d, 7e and 7f). These findings suggest that 3’UTR lengthening is negatively associated with translation activity in human NPCs, aligning with prior polysome profiling studies in HEK293T cells^50^.

## Discussion

To leverage the high coverage and accuracy of 3’ end information in LR RNA-seq data, we developed APALORD, an R package, to analyze APA regulation between two different conditions. APALORD identifies PAS peaks from LR RNA-seq data at single nucleotide resolution, quantifies the relative PAU for multiple PASs within a host gene, and compares the PAU change at the individual PAS level and APA change at the single-gene level between biological groups. Applying APALORD to direct RNA-seq data from hESCs and hESC-derived neurons, we identified PASs with comparable accuracy to PolyA_DB reference annotations, which are based on short read RNA-seq data generated from 3’ end tag-specific library preparation approach 3’READS. APA analysis of this direct RNA-seq dataset recapitulated a transcriptome-wide 3’UTR lengthening trend during neural differentiation. This trend correlated with an increased preference for PAS featuring the canonical upstream AAUAAA poly(A) signal and UGUA motif in neurons compared to hESCs. The AAUAAA hexamer sequence is the primary determinant of a strong PAS^53^, with upstream UGUA and downstream GU/U-rich elements^54^ further enhancing PAS strength^55^. Our findings align with prior investigations demonstrating that AAUAAA and UGUA defined PASs are enhanced in neurons^56^.

Longer 3’UTR isoforms, cleaved at distal PASs, are preferentially expressed in brain tissue and neurons, whereas shorter 3’UTR isoforms, cleaved at proximal PASs, are enriched in testis, liver and cancer cells^57,58,59,60,13^. Testis specific PASs exhibit a reduced prevalence of the canonical AAUAAA hexamer, contain unique DSEs, and produce shorter 3’UTRs^61,62^. In contrast, the distal PASs typically display strong consensus features of canonical PAS, including the AAUAAA hexamer and strong USEs and DSEs, and high evolutionary conservation^63,64^. It has been shown that the longer 3’UTR isoforms cleaved at the distal PASs can persist in the nucleoplasm, serving as precursors for further processing at weaker PASs^64^. Our analysis of the neuron-enriched PASs, independent of their relative position within the gene, reveals a higher incidence of strong consensus features compared to those enriched in proliferating hESCs. Collectively, these findings support a hypothesis that the global trend of 3’UTR lengthening in neurons arise from a nucleoplasmic blockade of further 3’ end processing at weaker PASs, favoring the retention of longer isoforms critical for neuronal function.

Oligo-dT-initiated reverse transcription (RT) for short-read RNA-seq often masks false 3’ ends caused by internal priming due to poor 3’ end resolution. However, in long-read RNA-seq using oligo-dT-initiated RT, internal priming generates truncated reads that are more pronounced and cannot be ignored. Applying APALORD to PacBio cDNA data, we quantified the prevalence of internal priming across genes and detected PASs. The relative position of internal priming-induced “PASs” to other “high-confidence” PASs in the gene body varies by gene, and gene-level APA change patterns are cell-, tissue-, or condition-specific, making their impact on APA analysis unpredictable. While internal priming reads contribute to the quantification of overall host gene expression, they cannot by simply excluded from downstream analysis. However, APALORD labels falsely recognized “PASs” affected by potential internal priming, enabling users to validate APA events with awareness of potential artifacts.

Taken together, APALORD offers a robust approach for identifying PASs from LR RNA-seq data and detecting APA changes across conditions. It provides precise 3’ end annotation independent of existing PAS references and quantifies both PAS-specific changes and cumulative 3’ end PAU shifts along the gene body. Unlike other APA methods, APALORD leverages the read counts of target PASs and genes within the dataset, enabling APA analysis even when biological replicates are not available or cost-prohibitive. Direct RNA-seq, which sequences the full length of the native RNA molecules without any transformation or modification, excels at capturing nucleotide modifications, 3’ polyA tail length and composition in addition to the messenger RNA sequence^65,66,67^. The growing interest in long-read RNA-seq for simultaneous analysis of differential gene and isoform expression, alternative splicing, APA, RNA modifications, and rare isoform discovery highlights its ability to explore the interconnected layers of mRNA processing. Building on this, APALORD is a powerful tool for APA research, leveraging LR RNA-seq data to precisely analyze APA events and unlock their role within the broader context of mRNA processing.

## Material and methods

### hES cell culture

H9-NGN2 hESC was generated by the UCONN Cell and Genome Engineering Core. Stem cells were grown for at least one passage on a 6-well tissue culture plate coated with Matrigel (Corning #CLS354277) in mTeSR Plus media (mTeSR+, StemCell Technologies #100-0276) over 3-4 days to ∼60-80% confluency. Cells were thoroughly singularized with 0.5 mL/well Accutase (StemCell Technologies #07920) for 9-10 mins at 37°C and resuspended by pipetting 15-20X with a P1000 in 1 mL mTeSR+/well. The cells were pooled and centrifuged in a 5 mL tube at room temperature in a swinging-bucket rotor at 220xg for 3 mins. The supernatant was removed from the cell pellet, which was resuspended in 1 mL Induction Medium (IM) [DMEM/F12 (ThermoFisher Scientific #11330032), 1x NEAA (ThermoFisher Scientific), 1x GlutaMAX10.7 (ThermoFisher Scientific), 66 μg/mL apo-transferrin (MilliporeSigma #T1147), 33 μg/mL insulin (MilliporeSigma #I6634), 66.7 μM putrescine (MilliporeSigma #5780), 13.4 nM progesterone (MilliporeSigma #P8783), and 20 nM sodium selenite (MilliporeSigma #S5261)] with 10 μM Y-27632 ROCK inhibitor (RI, Selleckchem #S1049). An aliquot was diluted 1/10 and counted in a Countess Cell Counter (ThermoFisher Scientific). For each well to be plated for induction, 200,000 cells, 2 mL of IM with RI and 2 mg/mL doxycycline (dox, MilliporeSigma #D5207) were plated on Matrigel plates, distributed evenly, and incubated overnight at 37°C. The remaining singularized cells were placed on ice, PBS was added to the tube to rinse, and these were centrifuged at 300xg for 3 mins. The resulting pellet was processed fresh from ice or snap-frozen in liquid nitrogen and stored at -80C for later processing as D0 samples. Media was replaced on day 1 and day 2 with fresh IM and dox. On day 3, cell wells were gently rinsed with PBS and singularized with Accutase as before, pooled and resuspended in 1 mL IM with RI. Cells were counted as before, and replicate wells were plated on 6-well poly-D-lysine- (MilliporeSigma #A-003-E) and laminin- (ThermoFisher Scientific #23017015) coated plates without dox at 500K/well in 2 mL IM with RI. On day 4, the media was fully and gently replaced with Cortical neuron Medium (CM) [½ DMEM/F12 and ½ Neurobasal media (ThermoFisher # 21103049) with 1x B27 supplement (ThermoFisher Scientific # 12587010), 1x Pen-Strep, 10 ng/mL BDNF (Peprotech # 450-02), 10 ng/mL GDNF (Peprotech #450-10), 10 ng/mL NT-3 (Peprotech #450-03), and 1 μg/mL laminin]. On day 7 induced neurons were collected by removing media, adding 1 mL ice-cold PBS per well, removing cells with a silicone cell scraper, and centrifuging each replicate at 300xg for 3 minutes in 1.5 mL tubes (in swinging bucket rotor). The resulting pellets were processed fresh from ice or snap-frozen in liquid nitrogen and stored at -80C for later processing as D7 samples.

### Nanopore direct RNA-seq library preparation and sequencing

Cell samples were subject to RNA extraction and on-column Dnase treatment using the PureLink RNA mini Kit (Invitrogen #12183025) coupled with PureLink Dnase (Invitrogen #12185010). DNA-free total RNA sample was used to prepare the library with the Nanopore direct RNA sequencing kit (Oxford Nanopore SQK-RNA004) per the manufacturer’s instruction. In brief, 2-3 ug of total RNA per sample was incubated with the RT adapter for 10-20 min at room temperature, followed by reverse transcription (RT) using Indora Reverse Transcriptase (New England Biolabs M0681L) and RNAClean XP beads (Beckman Coulter A63987). The eluted RT-RNA samples were then incubated with an RNA ligation adapter for 20-30 min at room temperature, followed by another RNAClean XP bead cleanup and then eluted in 15ul elution buffer. The libraries were quantified on Qubit using the dsDNA HS Assay Kit (Invitrogen Q32854). 40-70ng of the library was loaded onto a PromethION or MinION R10.4.1 flow cell and sequenced for up to 72 hours.

### LR RNA-seq data analysis

Dorado (Oxford Nanopore Technologies) was used to basecall the direct RNA-seq read sequences (sup@latest --min-qscore 0 --estimate-poly-a). Basecalled direct RNA reads or cDNA reads (from publicly available datasets) were aligned against the human genome (hg38) using Minimap2 (-ax splice -t 7 -B 3 -O 3,20 --junc-bed GENCODE v43 annotation). IsoQuant was then applied to each sample bam file individually to assign read to unique genes, using only primary alignments (--data_type nanopore -- gene_quantification unique_only --splice_correction_strategy default_ont -- no_secondary). The resulting output directories served as the input path for APALORD to load the data for downstream analysis. GO analysis was performed in R using clusterProfiler package^68^. R package BSgenome.Hsapiens.UCSC.hg38 was loaded in R to extract sequences around cleavage sites and PAS peaks according to the genome coordinates^69^.

## Data Availability

Direct RNA-seq data is deposited in SRA under BioProject PRJNA1247144. Encode PacBio LR RNA-seq data collected from dorsolateral prefrontal cortex (ENCSR205QMF, ENCSR094NFM, and ENCSR463IDK) and right lobe of liver (ENCSR657HJW and ENCSR293MOX) were used for APA analysis. Nanopore R2C2 LR RNA-seq data of human ESC and NPC polysome fractions analyzed in this work were downloaded through the Gene Expression Omnibus Short Read Archive (GSE244655).

## Code availability

Software is available at GitHub [https://github.com/markandtwin/APALORD].

## Acknowledgements

We thank Michael Guertin (University of Connecticut School of Medicine), and Jeremy Sanford (University of California, Santa Cruz) for insights and discussion on the manuscript. Thanks to Miura lab members for reading and providing input on the manuscript. This work was supported by NIGMS grant R35GM138319 awarded to P.M.

## Contributions

Conceptualization, Z.Z., and P.M; Methodology, Z.Z., D.S. and P.M.; Sample preparation, H.GD. and Z.Z., Data Analysis, Z.Z.; Package development, Z.Z., L.Z., D.S. and P.M.; Writing, Z.Z., and P.M.; Funding Acquisition, P.M.; Supervision, P.M.

---

**Supplementary Fig.1:**
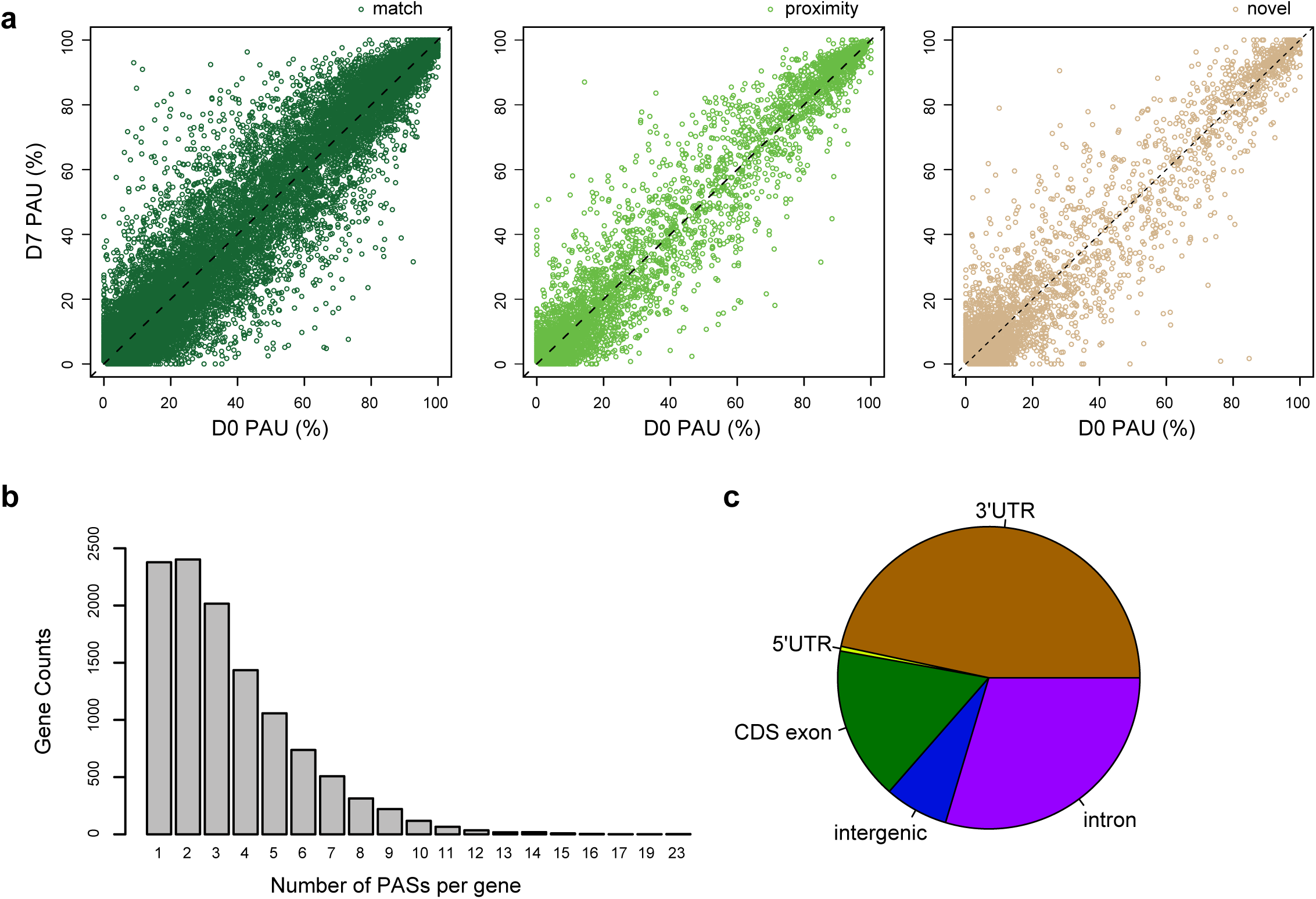
APALORD performance in PAS calling from direct RNA-seq of hESCs and hESC-derived neurons. **a-c** PAU data of match (**a**), proximity (**b**) or novel (**c**) PASs identified by APALORD in hESC (D0) and hESC-derived neuron (D7) samples. match: PASs aligned with PolyA_DB-annotated PASs; proximity: PASs within 20 nt of annotated PASs; novel: PASs outside of match and proximity criteria. **b** Number of genes with varying counts of PASs identified by APALORD in hESC (D0) and hESC-derived neuron (D7) samples. **c** Characterizing the gene body residence of novel PASs identified by APALORD in hESC (D0) and hESC-derived neuron (D7) samples.

**Supplementary Fig.2:**
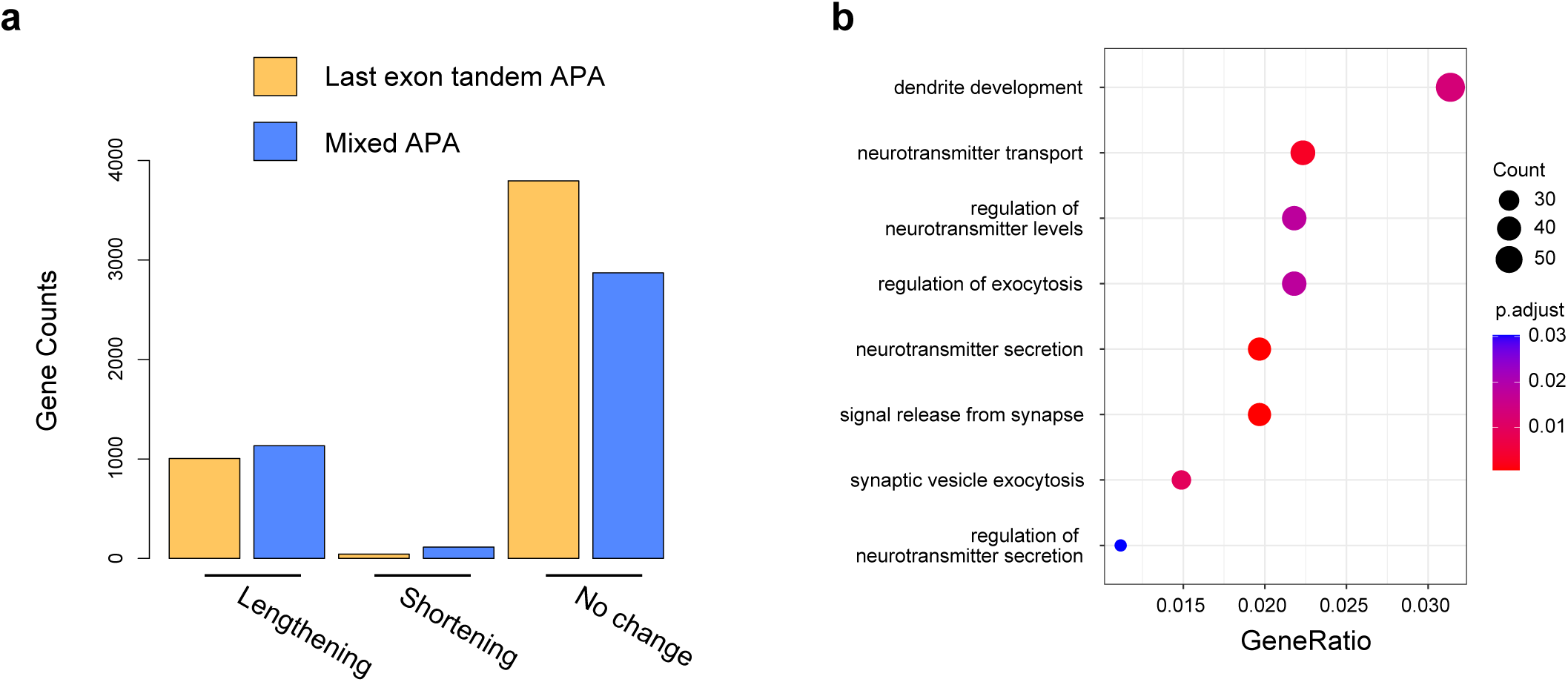
APA types and GO analysis of neural differentiation-regulated APA genes identified by APALORD. **a** Counts of last exon tandem APA and mixed APA genes identified by APALORD, categorized by 3’UTR change during neural differentiation (lengthening, shortening or no change) as defined in Fig.3d. **b** GO analysis of biological processes for genes with significant 3’UTR lengthening in hESC-derived neurons compared to hESCs.

**Supplementary Fig.3:**
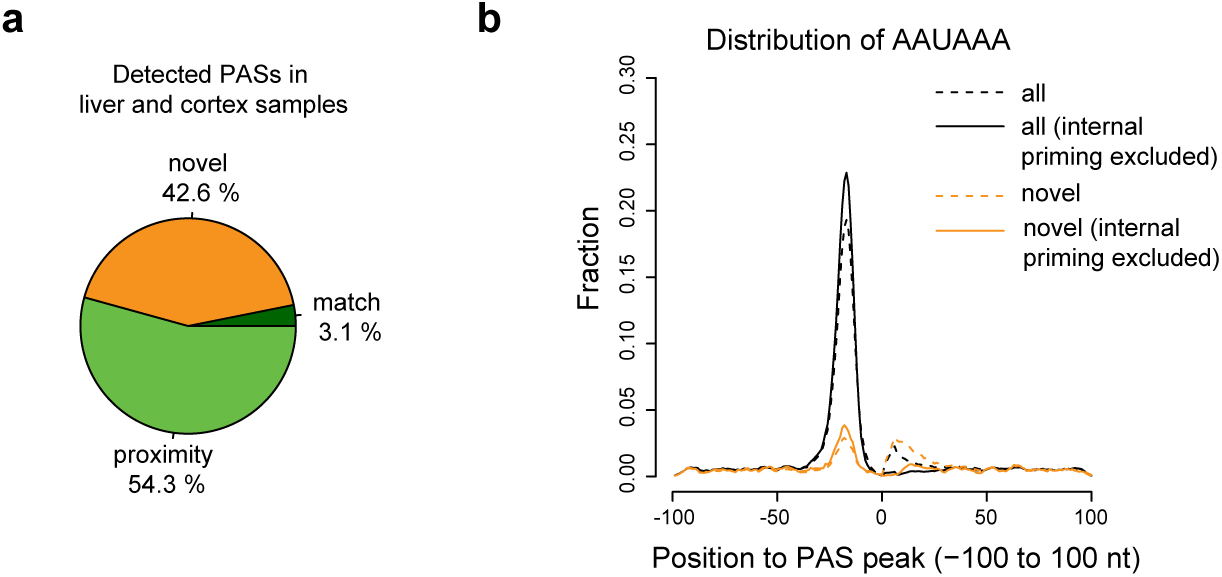
**a** All the PASs identified by APALORD (without internal priming filtering) from PacBio cDNA data of human cortex and liver, categorized as: match (aligned with annotated PASs in the PolyA_DB reference), proximity (within 20 nt of an annotated PAS), novel (outside match and proximity criteria). **b** Proportion of PASs with the canonical AAUAAA hexamer within the sequence surrounding the peak cleavage site (-100 nt to +100 nt), before or after internal priming filtering. Categories of PASs as defined in **a**.

**Supplementary Fig.4:**
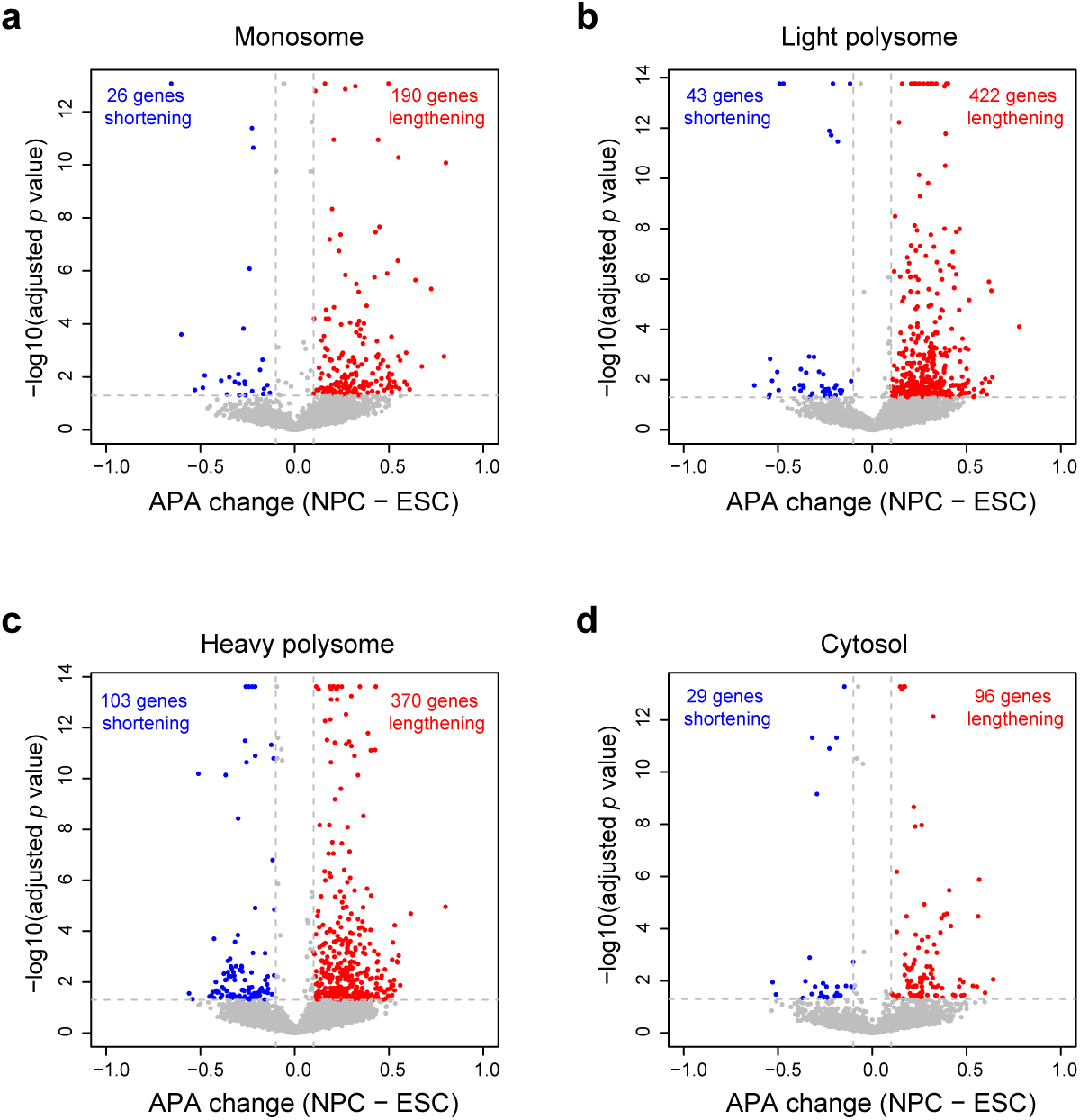
**a-d** Transcriptome-wide APA changes quantified by APALORD in NPCs compared ESCs across fractions: monosome polysome (**a**), light polysome (**b**) and heavy polysome (**c**) and cytosol (**d**).

